# Universal and taxon-specific trends in protein sequences as a function of age

**DOI:** 10.1101/2020.03.26.010728

**Authors:** Jennifer E James, Sara M Willis, Paul G Nelson, Catherine Weibel, Luke J Kosinski, Joanna Masel

**Affiliations:** Department of Ecology and Evolutionary Biology, University of Arizona, Tucson, Arizona 85721, USA; Department of Molecular Cell Biology, University of Arizona, Tucson, Arizona 85721, USA

## Abstract

Extant protein-coding sequences span a huge range of ages, from those that emerged only recently in particular lineages, to those present in the last universal common ancestor. Because evolution has had less time to act on young sequences, there might be “phylostratigraphy” trends in any properties that evolve slowly with age. Indeed, a long-term reduction in hydrophobicity and in hydrophobic clustering has been found in previous, taxonomically restricted studies. Here we perform integrated phylostratigraphy across 435 fully sequenced and dated eukaryotic species, using sensitive HMM methods to detect homology of protein domains (which may vary in age within the same gene), and applying a variety of quality filters. We find that the reduction in hydrophobic clustering is universal across diverse lineages, showing limited sign of saturation. But the tendency for young domains to have higher protein structural disorder, driven primarily by more hydrophilic amino acids, is found only among young animal domains, and not young plant domains, nor ancient domains predating the existence of the last eukaryotic common ancestor. Among ancient domains, trends in amino acid composition reflect the order of recruitment into the genetic code, suggesting that events during the earliest stages of life on earth continue to have an impact on the composition of ancient sequences.

## Introduction

New protein-coding genes emerge over time, either through a combination of rearrangement, duplication and divergence from ancestrally coding sequences, or *de novo* from previously non-coding DNA (for a review see (Van Oss & Carvunis 2019)). Although genes clearly originated *de novo* during the emergence of life, *de novo* gene birth after this unique event was once believed to be vanishingly rare (Jacob 1977; Keese & Gibbs 1992; Zuckerkandl 1975). Today, *de novo* gene birth is increasingly acknowledged to be real and important, with well-documented examples confirmed in diverse taxa (Baalsrud et al. 2018; Cai et al. 2008; Khalturin et al. 2009; Milde et al. 2009), despite the technical challenges associated with correctly identifying *de novo* genes (McLysaght & Hurst 2016). Compared to genes born early in the history of life, recently emerged *de novo* genes have had little time to evolve and adapt. If their properties converge only slowly to those of ancient genes, this creates trends in gene properties with age (Neme & Tautz 2013).

It has been claimed that younger genes encode shorter and faster evolving proteins (Lipman et al. 2002; Toll-Riera et al. 2012) that are more disordered, containing fewer hydrophobic amino acids (Wilson et al. 2017), which are more clustered together along the primary sequence (Foy et al. 2019). Foy et al. (2019) suggest that the two latter trends, in hydrophobicity and in its clustering, together indicate that evolution is slow to find the sophisticated folding strategies employed by older proteins, which minimize their propensity to form aggregates within cells despite relatively high overall hydrophobicity.

The universality of some of these trends has, however, been questioned. In particular, some authors have found that in some taxa, such as *Saccharomyces cerevisiae* (Carvunis et al. 2012; Vakirlis et al. 2018), young genes are in fact less disordered than older genes. Random sequences with high %GC tend to encode polypeptides with higher ISD (Ángyán et al. 2012). Differences in genomic GC content between species might therefore shape disorder at the time of gene birth differently (Basile et al. 2016; Van Oss & Carvunis 2019).

Alternatively, the low disorder of young “protogenes” may be an artifact of including sequences that do not meet the evolutionary definition of functionality, which requires that the loss of the gene be deleterious, i.e. have a selection coefficient of less than zero (Graur et al. 2013). In a species whose %GC gives rise to low structural disorder in non-functional protogenes, the pooling of high-disorder functional and low-disorder non-functional genes in varying proportions can create a spurious trend with age. Indeed, the direction of the disorder trend in *S. cerevisiae* was reversed when dubious gene candidates were removed from the data of Carvunis (2012) (Wilson et al. 2017). Nevertheless, with phylostratigraphy so far studying a limited set of relatively well-annotated focal species, it remains possible that trends in protein properties as a function of age are not universal, and instead depend on taxonomic group.

Identifying phylostratigraphy trends as a function of gene age requires accurate age estimates. Gene ages are based on the date of the most basal node in the phylogeny of lineages containing homologs (Domazet-Lošo et al. 2007). But when sequences are highly divergent, programs used to detect homologs, such as BLASTp (Altschul et al. 1990) are prone to false negatives, and thus underestimate gene age (Elhaik et al. 2006; McLysaght & Hurst 2016; Moyers & Zhang 2015; Moyers & Zhang 2017; Wolfe 2004). This is particularly problematic when studying protein properties such as length, evolutionary rate, and degree of conserved structure, because these properties themselves directly impact our ability to detect sequence similarity. Simulation studies have shown that these biases in homology detection have the potential to drive spurious phylostratigraphic trends (Elhaik et al. 2006; Moyers & Zhang 2015; Moyers & Zhang 2016). However, the number of ‘true’ young genes identified by phylostratigraphy studies may dwarf the numbers of genes that are falsely identified as young due to errors in homology detection, making that bias insufficient to explain previously observed trends (Domazet-Lošo et al. 2017). Statistically correcting for length and evolutionary rate did not remove observed trends in intrinsic structural disorder (Wilson et al. 2017) or clustering (Foy et al. 2019). While homology detection clearly has a false negative problem, many false negatives may be random errors, which merely reduce power in detecting trends, rather than systematic errors, which can create spurious trends.

Whatever the issues of random failure and bias, abandoning homology detection is not a viable option. Because homologs share an evolutionary history, they are not statistically independent. But many –omic studies treat genes as independent datapoints, and are thus flawed due to pseudoreplication. Phylostratigraphy can solve this problem by making each datapoint represent the independent evolutionary origin of a protein sequence, avoiding phylogenetic confounding (Felsenstein 1985; Thornton & DeSalle 2000). It is therefore critical that we improve upon the BLAST-based homology detection methods used within phylostratigraphy, rather than abandon the effort.

More recently developed methods, such as PSI-BLAST (Altschul & Koonin 1998) and HMMER3 (Finn et al. 2011), are more sensitive to distant homology, while still being computationally efficient. PSI-BLAST is a heuristic method that constructs a position-specific score matrix from the alignment of known homologs, which can then be used to iteratively search a protein database for additional homologs. HMMER instead constructs a profile HMM – a probabilistic model of the alignment – and is thus based on a formal statistical framework, yielding superior treatment of indels (Eddy 2011). Unfortunately, because both methods are based on iterative searches, both are vulnerable to model corruption, where a single false positive hit to a non-homologous sequence will pull in many more non-homologous sequences, potentially snowballing over multiple iterations (Pearson et al. 2017). Automated pipelines for whole-genome analysis are technically challenging, and HMMER is generally used with manual supervision of alignments.

Here we increase the sensitivity of homology detection during phylostratigraphy by using the manually-curated pfam database, which was constructed using a HMMER3 pipeline. For each pfam, an HMM was built from high quality seed alignments, homologs were then found, and matches above a curated threshold were re-aligned back to the HMM (El-gebali et al. 2019). Pfams are intended to correspond to protein domains, which are structural units, capable of folding independently (Holm & Sander 1994, for a review discussing domain definitions and identification, see Ponting & Russell 2002). These domains are often considered to be the ‘true’ units of homology, with full proteins made up of one or more domains in a variety of different combinations (Bagowski et al. 2010; Chothia et al. 2003; Koonin et al. 2000; Moore et al. 2008).

It is not known whether observed patterns as a function of apparent age get stronger or weaker when homology detection methods are improved. If most false negatives come from random error, improved methods might increase the signal to noise ratio and hence strengthen trends (McLysaght & Hurst 2016). But if systematic error drives observed trends in protein properties in real datasets, simulation studies suggest that reducing false negatives will reduce the strength of spurious trends driven by homology detection bias (Moyers & Zhang 2018).

Here we apply HMMER3-based phylostratigraphy to a broad range of taxa. A key innovation is to use pfam domains rather than BLASTp as our unit of homology. A second innovation is that instead of one or a few focal species, we have 435 fully sequenced species, each of which can be thought of as focal, allowing us to evaluate how general trends are across different phylogenetic groups. We use only those pfams that are present in at least two species, a quality filter that ensures the exclusion of non-genic contaminants from our dataset. We also analyse patterns of protein evolution in full genes, dating these by the oldest pfams that they contain, making results comparable to earlier phylostratigraphies based on the oldest short segment BLASTp hits (Albà & Castresana 2005; Domazet-Lošo et al. 2007). Additionally, to avoid spurious trends most likely to be due to homology detection bias, we avoid the protein metrics that most strongly affect homology detection (i.e. evolutionary rate and length), and focus on trends in properties expected *a priori* to have less effect on homology detection, such as the hydrophobicity and the degree of clustering of hydrophobic amino acid residues. Our large dataset gives us unprecedented power to detect trends in long-term evolution. It also enables us to detect finer-scale trends, such as changes in the frequencies of each of the 20 amino acids.

## Results

In this study, we analyzed all protein-coding genes containing at least one pfam annotated in the genomes of 435 species, with a total of 8,209 pfams identified (see Methods). Eukaryotic species were included in our dataset if they were marked “Complete” by GOLD (Mukherjee et al. 2019), and also present in TimeTree (Hedges et al. 2006), a phylogenetic database which we used to assign evolutionary ages to pfams. The genes in our dataset were then assigned ages based on the oldest pfam they contained. Pfams are likely to be the most highly conserved portions of genes, and thus adding BLASTp searches of other gene regions is not likely to push back the ages of many genes. We exclude genes without a pfam, which helps remove sequences that are not truly genic, but may underestimate disorder, especially in the youngest, most disordered genes that are least likely to have annotated domains. We also use a variety of quality filters to exclude pfams that may be due to contaminants, annotation errors, or horizontal gene transfer (see Methods). Where not otherwise indicated, we focus on non-transmembrane pfams. To avoid pseudoreplication caused by phylogenetic confounding, we treat the average across all homologs of a pfam or gene as a single data point in our analyses (see Methods).

The large size of our dataset gives us sufficient resolution to investigate trends in the prevalence of each of the 20 amino acids. Our dataset is also taxonomically varied, allowing us to address the question of whether previously reported trends are general across taxonomic groups.

### Trends in ISD

Young genes (Foy et al. 2019; Mukherjee et al. 2015; Willis & Masel 2018; Wilson et al. 2017) and domains (Bornberg-Bauer & Albà 2013; Buljan & Bateman 2009; Ekman & Elofsson 2010; Moore & Bornberg-Bauer 2012) have been reported to have high ISD, although some have claimed this depends on taxon (Vakirlis et al. 2018). We confirm that across our entire dataset, ISD is higher in young pfams (figure 1, linear model: R^2^ = 0.13, p = 3×10^−245^) and genes (Supplementary figure 1A, linear model for all genes: R^2^ = 0.057, p =1×10^−229^). Our improved methodology increased the steepness of the relationship between gene ISD and gene age in mouse genes above that previously estimated by Foy et al. (2019) (Supplementary Fig. 1B), from −0.028 to −0.050; note that mouse genes have a steeper slope than our results across all taxa.

**Figure 1.**
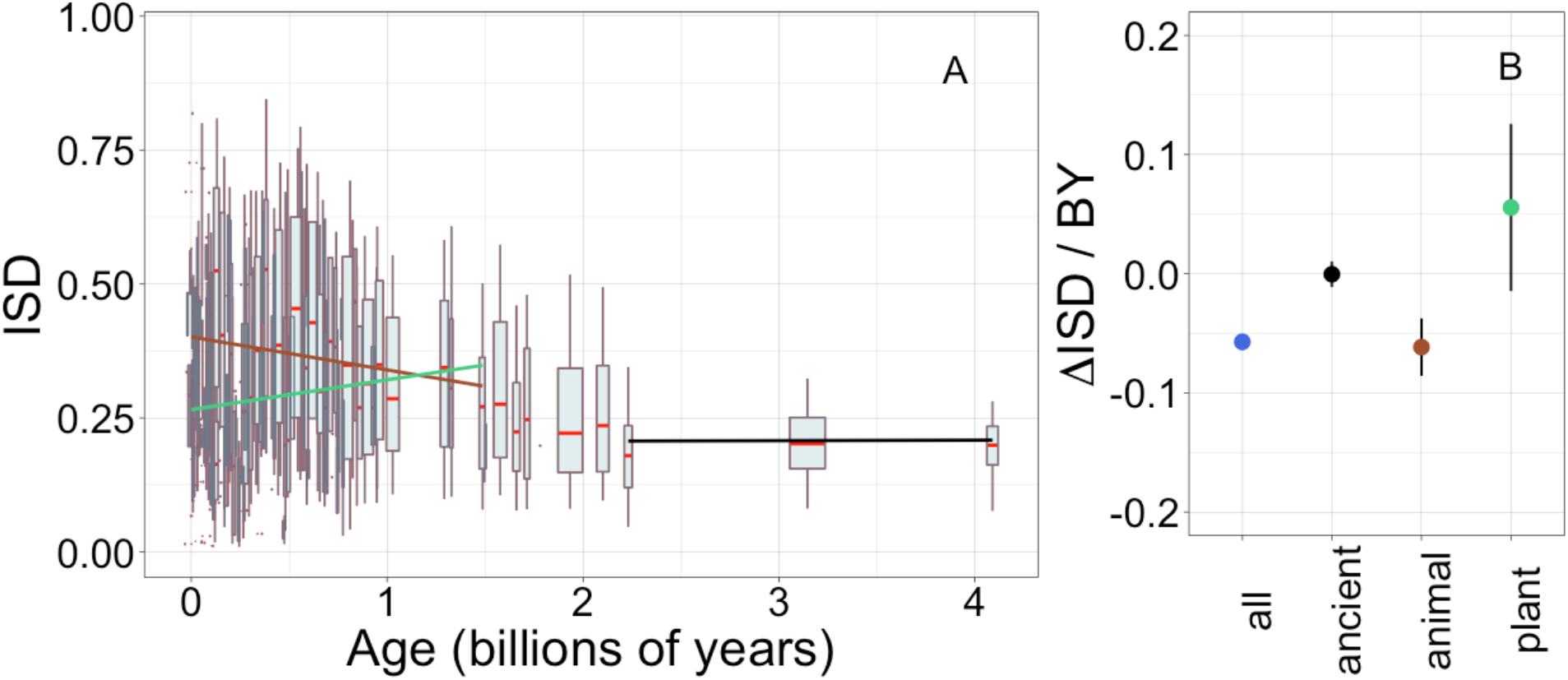
Young domains have high ISD, but this trend is driven exclusively by recent animal domains. Results are for non-transmembrane pfams. A) The brown linear regression was calculated for recent animal pfams (slope=−0.062, R^2^ = 0.0097, p = 6×10^−7^), green for recent plant pfams (slope=0.056, p = 0.1), and black line over ancient pfams in all lineages (slope = −4.2×10^−4^, p = 0.9). Slopes represent the decrease in average IUPred2 score, i.e. the predicted propensity of the average amino acid to be disordered, per billion years. Each data point consists of the average across all instances of homologous pfams, across all species in which it occurs. Phylostratigraphy assigns these to age classes, dated using timetree. Each age class is represented by a weighted box plot, where the width of the plot indicates the number of pfams in that age class. The median is shown in red, with the boxes representing upper and lower quartiles (the 75^th^ and 25^th^ percentile), and the whiskers indicating 9 and 91 quantiles. For age classes with only a single pfam, values are presented as small red dots. For clarity of presentation our plots do not show outliers. B) Phylostratigraphy slopes for pfams calculated over different subsets of the data are plotted with their 95% confidence intervals. The point colors correspond to the regression slopes in A).

The slope of the relationship between ISD and age is 1.8-fold steeper for pfams (Fig. 1A) than for full genes (Supplementary Fig. 1A); this is unsurprising, because whole genes are made up of combinations of sequences of different ages. The fact that protein domains are a more fundamental evolutionary unit than genes (Bornberg-Bauer et al. 2005; Moore et al. 2008), which is reflected in stronger relationships with pfam age, demonstrates an advantage of using a pfam-based phylostratigraphy approach. In results below, we will therefore focus on pfam domains, although we note that results are similar, but with smaller effect sizes, when calculated over genes.

ISD scores are similar among the oldest phylostrata. It is possible that pfams eventually reach an optimum level of ISD due to selection against over-stabilization (Daniel et al. 2003; DePristo et al. 2005; Fields 2001). Alternatively, there may be an optimal tradeoff point between having enough order to fold compactly while minimizing the risk of misfolding or aggregating (Bertram & Masel 2020).

To investigate how the ISD trend varies over time, we recalculated phylostratigraphy slopes over age-restricted subsets of our data. We also compared animal vs. plant lineages, given that these are the two kingdoms for which we have the most data (343 animal genomes and 87 plant genomes). The trend in ISD is not consistent (Fig. 1B), but is instead driven by recent animal pfams (in which we include all pfams that emerged after the divergence of animals/fungi from plants, 1496 MYA, that are now present in animals).

There is no significant change in ISD over “ancient” pfams (those that emerged prior to the last eukaryotic common ancestor (LECA), which is estimated to have existed around 2101 MYA), nor over recent plant pfams (i.e. all pfams found in plants that are younger than 1496 MY old). Note that we have relatively few plant-specific pfams in our dataset compared to animal-specific pfams. This is likely due to differences in genome quality and annotation between the groups, and the corresponding lack of available plant genomes that meet our quality standards. However, it is also possible that these numbers reflect a biological reality that animal proteomes are richer in domains (Ramirez-Sanchez et al. 2016).

Recent animal pfams have higher mean disorder (0.36) than recent plant pfams (0.26) (figure 1, Welch’s t-test p = 4×10^−5^), with the latter still higher than ancient pfams (0.21). The difference between animals and plants does not reflect differences in the birth process alone; even in ancient pfams that are shared by plants and animals, mean ISD is 0.27 in animals vs. 0.24 in plants (Wilcoxon signed rank test (a paired, non-parametric test) on difference p = 0.005). We conclude that animal domains experience more selection for higher ISD than plants do, with this difference being more pronounced in young, lineage-specific domains.

### Trends in amino acid composition

IUPred scoring of ISD primarily reflects amino acid composition, with hydrophilicity being a major determinant of disorder (Dosztányi, Csizmók, et al. 2005). Our larger dataset has sufficient power to investigate age trends in the frequencies of each of the 20 amino acids individually, rather than just trends in the single IUPred summary statistic. This can reveal which amino acids drive our ISD results, as well as other potentially interesting patterns in amino acid occurrence.

Trends in amino acid composition with age among ancient pfams are essentially identical whether they are assessed using only plant data or only animal data (Fig. 2A, Spearman’s rho = 0.94, p = 6×10^−6^, unweighted Pearson’s R = 0.98, p = 8×10^−14^). In subsequent analyses beyond Figure 2, we therefore pool data across all species in our dataset to calculate the phylostratigraphy slopes among ancient pfams with higher resolution.

**Figure 2.**
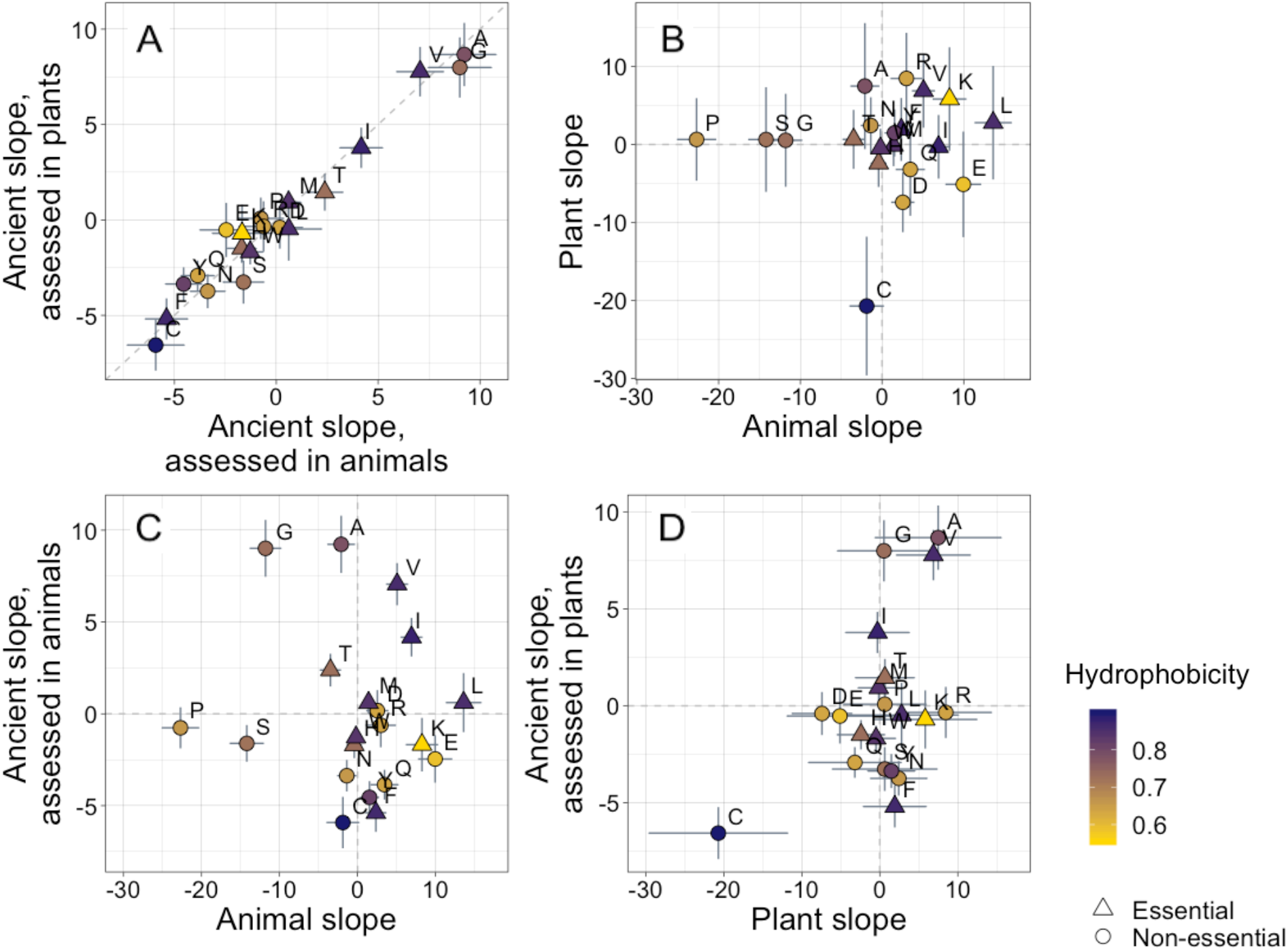
Trends in amino acid composition as a function of age differ across lineages. Results are shown for non-transmembrane pfams. Phylostratigraphy slopes are in units of the change in percentage points of an amino acid per billion years. ‘Ancient’ refers to pfams older than 2101 MY, assessed in only plant or only animal instances, whereas ‘plant’ and ‘animal’ slopes are calculated over pfams appearing after the divergence between the animal/fungi and plant lineages, 1496 MY, again assessed in only plant or only animal instances. Lines indicate the standard errors on the slopes. Points are color-coded by their hydrophobicity, as measured by 1-mean relative solvent accessibility (RSA) (Tien et al. 2013), such that buried, hydrophobic amino acids are dark purple, and exposed, hydrophilic amino acids are yellow. RSA scores are based on the water-accessible surface area of amino acids in a training set of proteins of known structure. Amino acid shapes indicate whether they are essential in animals. In A), y=x is shown as a dashed line, in all other plots dashed lines for x=0 and y=0 are shown for clarity. In A), Spearman’s rho = 0.94, p = 6×10^−6^. Spearman correlations for B), C) and D) are not significant (p = 0.8, 0.8, and 0.2, respectively).

In contrast, recent animal pfams and recent plant pfams show different trends in amino acid composition with age, with slopes of similar magnitude (Welch’s t-test on absolute slope values p = 0.3), but no correlation in value (Fig 2B, p = 0.8). Recent composition trends are mostly unrelated to ancient trends, with no relationship for animals (Fig. 2C, p = 0.8), and a weak correlation for plants that is not significant (Fig. 2D, p = 0.2) and is entirely driven by cysteine.

In plants, cysteine has the steepest phylostratigraphy slope. Miseta & Csutora (2000) previously reported that %cysteine content decreased with “complexity”, from mammals, to plants, to a set of single celled organisms that included both photosynthesizing and non-photosynthesizing bacteria and archaea. Over all pfams, our results agree with the findings of Miseta and Csutora, in that cysteine is slightly more common in animals (2.5%) than in plants (2.2%). But surprisingly, young, plant-specific pfams have 3.5% cysteine, as opposed to 2.6% in animal-specific pfams, and 2.1% in ancient pfams.

Photosynthesis likely creates greater oxidative stress in plants than in animals, which is likely to disrupt disulfide bonds. But in the training data used by IUPred, cysteine residues are mostly in stable disulfide bonds (Mészáros et al. 2018), so IUPred scores cysteines as highly order promoting. If unpaired cysteines are common in plants, this might obscure a trend in ISD. However, recalculating IUPred scores from sequences from which we had excised cysteine resulted in very little change in our ISD results (Supplementary Fig. 2).

It is possible that young plant domains are cysteine-rich because cysteine is relatively available in the cell. Producing cysteine is the final step in the assimilation of sulfur by plants. Additionally, cysteine production enables the detoxification of reactive oxygen species produced during photosynthesis, and is essential to chloroplast function (Bermúdez et al. 2010). In the cytosol, levels of cysteine appear to play a role in maintaining redox capacity, and in pathogen resistance (Romero et al. 2014). But however abundant cysteine is in the cytosol, its incorporation into protein sequences, while inexpensive in terms of raw materials, might be deleterious in terms of risk of oxidative damage and unstable disulfide bonds. This could result in natural selection removing cysteines from plant protein sequences over time.

After cysteine, the two amino acids that are most enriched in young plant pfams are glutamic acid and aspartic acid, E and D. These are the most abundant amino acids in plants as a whole (Kumar et al. 2017), and may therefore also be enriched in young plant domains due to their high availability.

In young animal pfams, the two amino acids that are most enriched are serine and proline. These are two of the amino acids previously identified as most responsible for differences between eukaryotes and prokaryotes via the greater quantity of ‘linker’ regions (protein sequences between pfams) of eukaryotic proteins (Basile et al. 2019). This suggests that the signal in eukaryotic linkers reflects an animal-specific trend across all recent protein-coding sequences, both when they are annotated as domains and when they are not. This is perhaps unsurprising; the simple categorization of sequences as either ‘pfam’ or ‘linker’ is artificial, and might introduce a variety of biases.

Because animal species with more tyrosine kinases have less tyrosine, it has been argued that selection against deleterious tyrosine phosphorylation has driven a decline in tyrosine content in metazoa (Tan et al. 2009). However, evidence that selection drives tyrosine loss is limited (Pandya et al. 2015), and it has been argued that changes in GC content might explain trends in tyrosine (Su et al. 2011, although see Tan et al. 2011). Examining this question with a phylostratigraphy approach for the first time, we do not see evidence for tyrosine loss in animals; tyrosine has a relatively shallow phylostratigraphy slope that is indistinguishable from 0 (supplementary table 1). This agrees with the findings of Pandya et al. (2015), who found no significant difference between the frequency spectra of alleles that removed or created tyrosine, suggesting that tyrosine is neither strongly selected for nor against, with trends in tyrosine being modest compared with those of other amino acids. There is, however, more tyrosine in younger ancient pfams than in the oldest ancient pfams, suggesting an ancient rather than recent process of tyrosine loss (see supplementary table 1).

To assess whether animal-specific trends also reflect amino acid availability, we examined essential amino acids (i.e. amino acids that plants but not animals can synthesize (Guedes et al. 2011)), expecting them to be rarer in young animal domains. While this is the case (with 6 of 9 of their phylostratigraphy slopes being positive, Fig 2), the difference is not significant (Binomial test with 50% chance of a positive slope, one-tailed p = 0.3).

Instead, the ISD score returned by IUPred seems to best summarize animal-specific trends in amino acids. Metrics aimed more strictly at hydrophobicity, such as relative solvent accessibility (RSA) (Tien et al. 2013), do less well. While we do see a tendency for hydrophobic amino acids to have positive phylostratigraphy slopes among animal pfams and negative slopes among plant pfams (amino acids are colored by RSA in Figures 2 and 3), this correlation between RSA and phylostratigraphy slope is not significant for any of the lineage- or phylostrata-specific subsets of the dataset shown in Figure 2 (see supplementary fig. 3).

**Fig 3.**
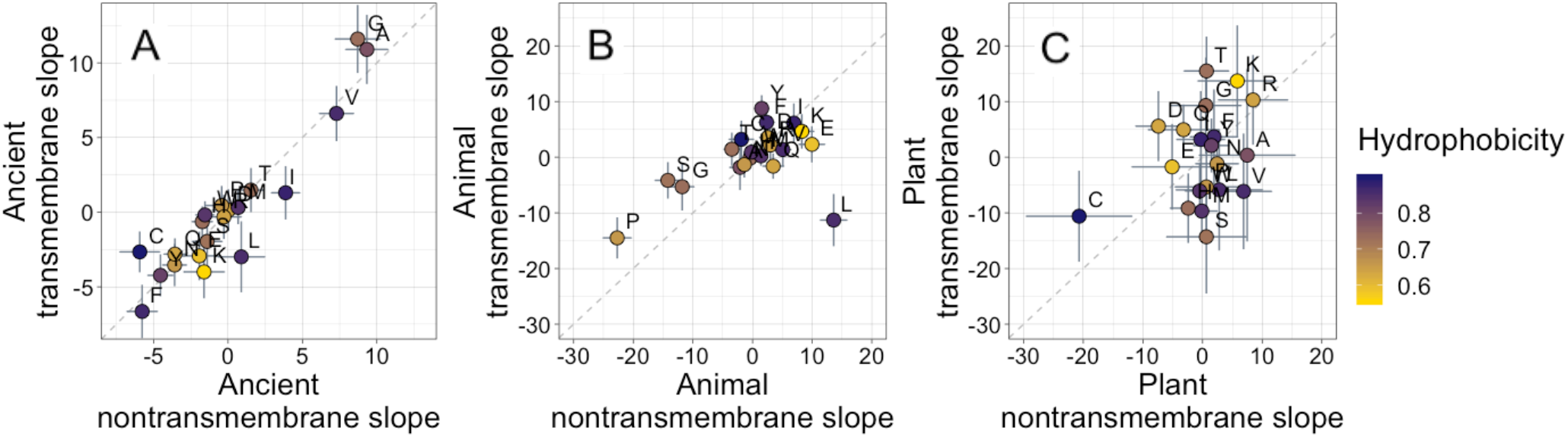
Ancient domains exhibit strikingly similar amino acid trends, whether transmembrane or non-transmembrane. Phylostratigraphy slopes are in units of percentage point change in composition per billion years. Taxonomic and temporal subsets of the data are as the same as in Figure 2. Lines indicate the standard errors on the slopes. Points are color-coded by their hydrophobicity, as measured by 1-RSA (Tien et al. 2013), such that buried, hydrophobic amino acids are dark blue, and exposed, hydrophilic amino acids are yellow. In all plots, y=x is shown as a dashed line. Congruence is strong for ancient pfams in A) (Spearman’s rho = 0.83, p = 5×10^−6^), weakly significant for animal-specific pfams in B) (p = 0.06) where transmembrane trends tend to be weaker and leucine is an outlier, and not detectable in our weakly powered plant-specific data set in C) (p = 0.3).

What is more, if hydrophobicity were the main determinant of phylostratigraphy slopes, we would expect amino acid composition to evolve differently in a hydrophobic membrane environment than in the cytosol. However, amino acid slopes are highly correlated between transmembrane and non-transmembrane ancient pfams (Figure 3A). Clearly, trends among ancient domains are not primarily driven by hydrophobicity. Amino acid slopes are also weakly, albeit not significantly, correlated between transmembrane and non-transmembrane pfams in animals (Figure 3B). Breaking this animal correlation is leucine, a clear outlier in that it is enriched in young transmembrane pfams, but depleted in young non-transmembrane pfams. Leucine also shows the same pattern to some degree among ancient pfams. There is very little power to detect a correlation in plants, should it exist (Figure 3C).

There has been concern that phylostratigraphy slopes could be driven by homology detection bias (Moyers & Zhang 2015; Moyers & Zhang 2016; Wilson et al. 2017). In this case, amino acids that are more changeable (Tourasse & Li 2000), making homology more difficult to detect, should be over-represented in young genes. However, we find if anything the opposite: more changeable amino acids are (slightly) enriched rather than depleted among the oldest of ancient pfams (Supplementary fig. 4, Spearman’s rho = 0.50, p = 0.02 for nontransmembrane pfams, and rho = 0.40, p = 0.08 for transmembrane ancient pfams). Our results are therefore not driven by homology detection bias even at extremely long timescales. There is no correlation between recent animal (p = 0.57) or recent plant (p = 0.7) phylostratigraphy slopes and amino acid changeability, and therefore no evidence that these results are affected by homology detection bias either. Furthermore, we note that homology detection bias is expected to create similar patterns for all taxa. The fact that the strength and even direction of amino acid trends can be different for different taxa is itself also evidence against a strong role for homology detection bias.

Given the striking similarity in ancient trends between transmembrane and non-transmembrane domains, we note that Jordan et al. (2005) claim ongoing trends related to the origin of the genetic code. Specifically, they claim that the amino acids incorporated earliest into the genetic code (Trifonov 2000) tend to be lost, while more recently acquired amino acids tend to be gained. Jordan et al. (2005) used a parsimony-based method of ancestral sequence reconstruction to detect trends in the amino acid composition of existing protein-coding sequences over time. The method has, however, been much criticized (Goldstein & Pollock 2006; Hurst et al. 2006; McDonald 2006).

While our amino acid phylostratigraphy slopes do not correlate with the flux values of Jordan et al. (2005) (Supplementary fig. 5), we nevertheless confirm the broader hypothesis by finding that among ancient domains, the very oldest domains are enriched in the amino acids that were recruited into the genome first, both in nontransmembrane domains (Spearman’s rho = −0.62, p = 0.004), and in transmembrane domains (Spearman’s rho = −0.55, p = 0.01). The order of recruitment ranking is taken from Trifonov (2000), who calculated a consensus chronology from the expectations of 40 amino acid ranking criteria, including such diverse factors as results from experiments into primordial conditions, amino acid complexity and thermostability. However, 3 of the 40 criteria were based on the amino acid compositions of protein assemblages. Therefore, to ensure non-circularity, we recalculated the consensus order, excluding these criteria and calculating the mean rank, as in Trifonov (2000) prior to their proposed second step of smoothing by a filtering procedure. This recalculation had very little effect on the order of amino acids, and our correlation in ancient nontransmembrane pfams remains significant (Fig. 4A, Spearman’s rho = −0.58, p = 0.008), although the correlation is no longer robust to multiple testing for ancient transmembrane pfams (Fig. 4B, Spearman’s rho = −0.47, p = 0.04).

**Fig 4.**
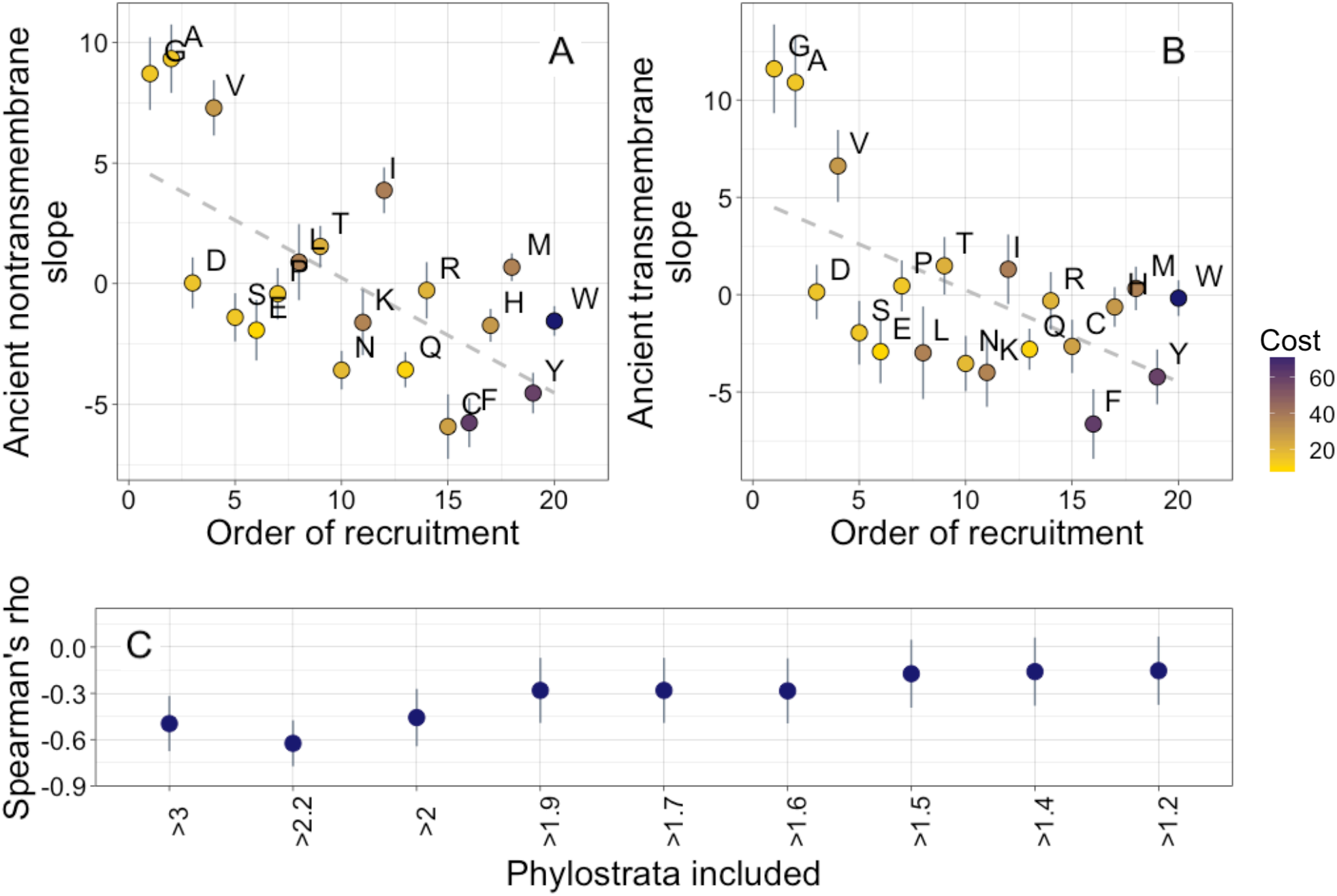
Ancient amino acid phylostratigraphy slopes reflect the order of recruitment of amino acids into the genetic code. Phylostratigraphy slopes for non-transmembrane (A) and transmembrane (B) pfams are in units of percentage points of composition per billion years, with lines indicating the standard errors on the slopes. Phylostratigraphy slopes for ancient pfams are calculated over all lineages, and include all pfams over 2101 MY. Consensus order is modified from Trifonov (2000) (see description in Results), and is given as rank data, such that the first amino acid to be recruited is given a rank of 1. Regression slopes are shown as dashed lines. Late-recruited amino acids are rare in the most ancient non-transmembrane pfams (A) Spearman’s rho = −0.58, p = 0.008 and transmembrane pfams (B), Spearman’s rho = −0.47, p = 0.04. Points are color-coded by their metabolic costliness to produce (measured as the number of high energy phosphate bonds required for synthesis, plus the energy lost due to the precursors used in the synthesis), as estimated for aerobic conditions in yeast (Raiford et al. 2008), such that the most costly amino acids are blue, and the least costly are yellow. C) The correlation (Spearman’s rho) between amino acid phylostratigraphy slope and order of recruitment over different subsets of our dataset. X-axis labels indicate the minimum age of pfams included, in billion years. Lines are the standard errors of the Spearman’s rho values, calculated using the Fisher transformation (Fisher 1915).

In contrast, the order of recruitment is not significantly correlated with phylostratigraphy slopes in more recent taxa (Supplementary fig. 6). This is in agreement with the broader hypothesis of Jordan et al. (2005), but hard to reconcile with alternative explanations for the correlation with order of amino acid recruitment that invoke a third correlated factor, such as relative metabolic costs.

Our interpretation of these results is that amino acids that were introduced late into the genetic code remained relatively rare for a prolonged period of time, well after the genetic code was complete. To determine for how long, we ask when the correlation between phylostratigraphy slope and order of recruitment is strongest (Fig. 4c). The strongest correlation is with slope over the three oldest phylostrata in our dataset. These span the pfams found in LUCA, through to pfams present in all extant eukaryotic but no prokaryotic lineages. The strength and significance of the correlation between order of recruitment and phylostratigraphy slope decreases if younger phylostrata are included, suggesting that from the last eukaryotic common ancestor (LECA) onward, the order of recruitment into the genetic code no longer shaped amino acid availability.

We do not believe our order of recruitment results to be a byproduct of a correlation with amino acid costliness. This is because there is no more than marginal significance for correlations between phylostratigraphy slope and the metabolic cost of amino acid production in yeast (Supplementary fig. 7; amino acids are color-coded by costliness in figure 4) (Raiford et al. 2008).

### Trends in Clustering

Finally, we consider whether there are any trends in amino acid order. Specifically, a temporal trend in the value of a ‘clustering’ metric has previously been reported for mouse gene families (Foy et al. 2019). This clustering metric calculates the degree to which hydrophobic amino acids tend to lie close together along the primary sequence, normalized to ensure that values do not depend on length or overall hydrophobicity (Irback et al. 1996). We confirm, using our considerably larger dataset, that young pfams (Fig 5A, slope = −0.037, R^2^ = 0.028, p = 1×10^−50^), and young proteins (Supplementary fig. 8A; slope= −0.031, R^2^ = 0.033, p = 5×10^−132^) have more clustering of their hydrophobic amino acids. In Supplementary fig. 8B, we show that our improved methods for assigning gene age result in a steeper slope for mouse genes than that reported by Foy et al. (2019): −0.056 instead of −0.045.

**Figure 5.**
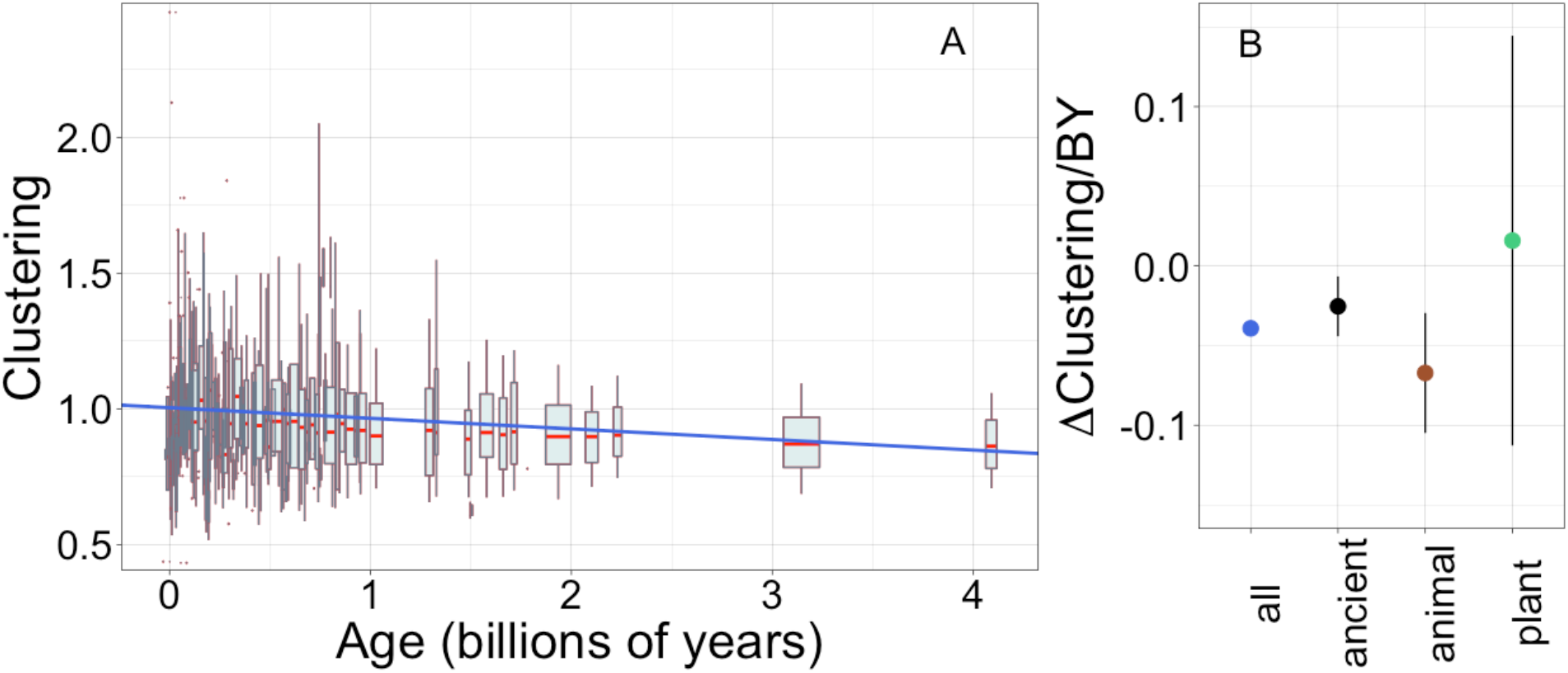
Young domains have more clustered hydrophobic amino acids (A), and the trend in clustering with age is consistent across time and taxonomic groups (B). Clustering has an expected value of 1 for randomly distributed amino acids. Results are shown for non-transmembrane pfams. A) Each data point consists of the average across all instances of homologous pfams, across all species in which it occurs. Phylostratigraphy assigns these to age classes, dated using timetree. Each age class is represented by a weighted box plot, where the width of the plot indicates the number of pfams in that age class. The median is shown in red, with the boxes representing upper and lower quartiles (the 75^th^ and 25^th^ percentile), and the whiskers indicating 9 and 91 quantiles. For age classes with only a single pfam, values are presented as red dots. For clarity of presentation our plots do not show outliers. The blue line is the linear regression slope calculated over all lineages. Slope= −0.037, R^2^ = 0.028, p = 1×10^−50^. Slope represents the average decrease in clustering score per billion years. B) Phylostratigraphy slopes (in units of change in clustering per billion years) for pfams are calculated over different phylostrata subsets of the data, and shown with their 95% confidence intervals. The left-most point corresponds to the regression slope in A).

Unlike the trend in ISD, the trend in clustering with age is remarkably consistent, both across lineages and age categories (figure 5B). While the confidence intervals increase for subsets that contain less data (such as the set of young plant pfams), they always clearly overlap the clustering slope calculated over all lineages and all phyostrata. This suggests that trends in clustering may be universal, with the dispersion of hydrophobic amino acids occurring at an approximately constant rate over the whole of evolutionary time. The data is also compatible with the trend slowing down, but not completely saturating, as it draws closer to an optimum in ancient pfams.

## Discussion

We have shown that there is a universal trend for young genes and pfam domains to have clustered hydrophobic amino acid residues, while old domains and genes have more evenly interspersed hydrophobic amino acids. In contrast, trends in amino acid content depend on taxon. Only animal trends are driven by young protein-coding sequences having high intrinsic structural disorder. Ancient trends, from LUCA to the last eukaryotic common ancestor LECA, are driven by the order of recruitment of amino acids into the genetic code. We have less power to decipher plant trends, but results are compatible with young plant sequences preferring easily available amino acids, with a particularly strong excess of cysteine, especially relative to the generally low levels at which abundant cellular cysteine is incorporated into plant proteins as a whole.

Our analysis relies on homology detection of pfam domains. It is important to consider whether systematic errors in homology detection could bias our results. Our improved hmmer-based homology detection methodology should reduce the false negative rate, and thus the scope for homology detection bias. But both our ISD and our clustering trends are stronger rather than weaker than those previously reported using blastp-based methods (Foy et al. 2019; Wilson et al. 2017). Homology detection bias also cannot explain why trends in ISD or amino acid composition are lineage-specific, nor the absence of correlation with amino acids’ evolutionary changeabilities. Additionally, old sequences are expected to be most affected by homology detection bias (Moyers & Zhang 2017), but it is more recent animal domains that drive the ISD result, which is *a priori* the trend most likely to be driven by homology detection bias. Overall, we find substantial evidence contradicting the suggestion that homology detection bias drives trends.

The rejection of homology detection bias does not imply that distant homology is always successfully detected, but rather that errors in age assignment might be primarily random rather than systematic. Evidence is accumulating that *de novo* gene birth is not, as once thought, vanishingly rare (or even non-existent) after the origin of life. A recent analysis exploited synteny to find that most taxonomically restricted genes are not the product of divergence, but have instead emerged *de novo* (Vakirlis et al. 2019). Another recent study excluded genes with insufficient power to detect ancient homology, and similarly found that evidence for *de novo* origin persisted in a substantial number of cases (Weisman et al. 2020). To get correct trends given random failures in homology detection, all we need is for apparent ages to be correlated with true ages.

Sequences can also be assigned as older than they really are, when they are incorrectly classified as present in distantly related species (false positives), due either to overly permissive homology detection cutoffs, or to contamination (see, for example, Longo et al. 2011; Merchant et al. 2014). In addition, some domains may be present in species due to horizontal gene transfer, which is problematic for the purpose of our analysis as it means that our annotation based on presence in a species does not reflect the evolutionary history of that domain (Liebeskind et al. 2016; Ravenhall et al. 2015). We used a number of filters to remove false positives and likely products of HGT from our analysis (see Methods). False positives are not expected to be subject to systematic error with respect to ISD or clustering (Moyers & Zhang 2018). In agreement with this, we found that these steps reduced the amount of ‘noise’ in our dataset, making the trends we observed stronger with each improved iteration of our quality control.

Our results are consistent with the hypothesis that selection acts to reduce the aggregation propensity of proteins. Polypeptide chains are intrinsically prone to forming aggregates that are toxic to cells (Chiti & Dobson 2017). Young animal genes avoid aggregation by being on average more disordered and hydrophilic (Wilson et al. 2017), with their few hydrophobic residues tending to cluster together along the protein chain (Foy et al. 2019). However, in order to be retained, sequences must also consistently serve a function within their host cells; this often requires them to fold, which is promoted by higher hydrophobicity. Frustration between these two aims might be resolved by sequences evolving more sophisticated strategies for folding while avoiding aggregation.

One such strategy appears to be the ordering of amino acid residues (Bertram & Masel 2020; Foy et al. 2019), with older, more evolved sequences having more evenly dispersed hydrophobic amino acid residues. Remarkably, we find that the trend in clustering is universal, even though the trend in disorder/hydrophobicity is restricted to animals. It is possible that pfam domains reach an optimal level of disorder after approximately 2000 million years, perhaps because over-stability is harmful (DePristo et al. 2005), or because a balance between folding and misfolding is reached. However, evolution does not appear to achieve such an optimum level of clustering. On all timescales, selection seems to act to decrease the clustering of hydrophobic amino acids.

Disordered regions can contain “sticky” amino acids at protein-protein interaction surfaces (Levy et al. 2012), whose hydrophobicity within a hydrophilic region will decrease clustering. More abundant proteins have fewer sticky residues within their disordered regions (Dubreuil et al. 2019). Older domains generally have higher protein abundance (Carvunis et al. 2012), which should cause lower stickiness and higher clustering, not the lower clustering seen in our results. Our clustering trend is therefore not driven by trends in protein “stickiness”, which is a possible contributor to aggregation propensity.

Protein length is another hidden factor that might contribute to the trends we see. It is *a priori* likely that younger proteins are shorter (Van Oss & Carvunis 2019) with a correspondingly higher surface to volume ratio, and thus a proportionately smaller hydrophobic core and higher ISD score. However, this cannot explain our results, because while younger animal domains do tend to be shorter, shorter animal domains are actually less disordered (R^2^ = 0.0025, p = 0.007). The reasons for this are unclear; length and disorder are, as expected from surface to volume considerations, negatively correlated in ancient domains and in recent plant domains.

Differences between taxa are surprising, with young plant domains having different amino acid compositions from young animal domains, and not sharing their preference for high ISD. Plant domains as a whole are more ordered, with ancient domains in plants also having lower ISD in plants than the same domains in animals (although the difference in ISD is most pronounced in lineage-specific domains). We speculate that plants are simply better at coping with proteostasis than other taxonomic groups, using mechanisms other than high ISD to protect themselves from protein misfolding and aggregation. Plant proteomes are rich in predicted amyloidogenic regions (Antonets & Nizhnikov 2017a), and yet there is a paucity of evidence for aggregate formation in plants under normal conditions (Antonets & Nizhnikov 2017b). This is perhaps due in part to the presence of molecules such as polyphenols, which inhibit aggregation (Velander et al. 2017). The effectiveness of plant molecules at inhibiting aggregation has also been related to plant longevity (Mohammad-Beigi et al. 2019). Young, ordered plant proteins may therefore be under weaker selection against aggregation than the same protein would be in an animal. Our dataset does not have enough plant genomes to give us power to detect whether selection acts on clustering in plants.

Our data are consistent with a hypothesis of selection acting in the usual way, i.e. that proteins born *de novo* go on to discover, through mutation, more sophisticated strategies for folding while avoiding aggregation and/or misfolding. A second, not mutually exclusive possibility is that the trends we observe are due to differential retention of pfam domains and genes, rather than descent with modification. Under this hypothesis, newborn sequences are diverse, and sequences with favored properties are lost less often during subsequent long-term evolution. The relative contribution of these two levels of selection could be investigated if quality control could be made stringent enough to obtain quantitatively reliable loss rates and diversification rates for pfam domains.

In conclusion, trends in the evolution of amino acid composition show surprising differences between taxonomic groups, and over different spans of time. Amino acid availability at time of *de novo* birth has a strong effect on domain composition. Even after billions of years of evolution, ancient domains remain influenced by which amino acids were more abundant as the genetic code was being formed. *De novo* birth and subsequent trends in plants may also be shaped by amino acid availabilities. In animals, amino acid composition seems to reflect young domains’ preference for high intrinsic structural disorder. However, protein sequences show a universal trend toward reducing their levels of hydrophobic clustering, no matter their amino acid composition, presumably in an attempt to find a balance between folding and misfolding. The only claim in evolutionary biology that is close to comparable in scope is the increase in body size in some taxa over the 3.5-billion-year history of life (Cope 1885; Heim et al. 2015; Payne et al. 2009).

## Materials and Methods

### Data compilation

We compiled a dataset of whole-genome sequences with a sequencing status of ‘complete’ from the GOLD database (Mukherjee et al. 2019, accessed August 7, 2018), resulting in a list of 1138 unique species. We then downloaded species from the Ensembl Biomart interface databases (Kinsella et al. 2011; Zerbino et al. 2018, fungi V40, plant V40, metazoan V40, protest v40 and main V93, accessed between 07/31/2018 - 09/12/2018). Of the resulting 306 species, 19 were not found in the GOLD species list and thus were excluded. We next searched the NCBI RefSeq repository (O’Leary et al. 2016, accessed between 09/27/2018 - 10/26/2018) for the remaining GOLD species, excluding archaeal, bacterial and viral genomes due to lack of phylogenetic resolution. Of the resulting 948 species, 344 had been annotated, were in the GOLD species list, and were not also in Ensembl Biomart.

We further required all species in our dataset to be present in TimeTree (Hedges et al. 2006), eliminating 163. Duplicates were removed (7 species), as were species using an alternative coding alphabet (2 species), or species for which there were genome annotation quality issues (such that we were able to identify unusually many more pfams from blast searches of intergenic regions than were present in the currently annotated genome, and/or encountered technical problems such as many annotated genes being rich in stop codons) (4 species), resulting in a final dataset of coding sequences of 455 unique species.

Protist genomes were retained, but used only as outgroups for dating (for a review, see Eme et al. 2014), while full protein properties were calculated for a core set of 435 animal, plant and fungal species. Protein-coding sequences of the 455 species in the curated list were then downloaded from the previously listed Ensembl repositories from their respective FTP sites, or from NCBI’s FTP RefSeq repository. For a full set of species used in the analysis, see Supplementary Table 2. Following genome acquisition, mitochondrial and chloroplast genes were removed from the dataset.

Finally, genes often have multiple annotated transcripts/proteins, resulting in non-independent datapoints. We chose a single transcript to represent each gene, specifically the closest homolog to the most closely related sister species in our dataset, identified using reciprocal Blastp (Altschul et al. 1990). If a gene failed to produce any hits below an e-value cutoff of 10^−3^, the longest gene transcript was chosen instead.

### Domain Annotation

Ensembl provided Pfam annotations for all protein transcripts, which are based on InterProScan with default parameters. These annotations were manually downloaded from the BioMart web interface and paired with their corresponding protein using Ensembl’s unique transcript identifiers. To make our dataset internally consistent, we replicated Ensembl’s annotation methodology by processing all NCBI sequences with InterProScan (Jones et al. 2014, v.5.20-69.0 downloaded June 20, 2018) using default parameters. All sequences without an associated Pfam were discarded from our analyses.

### Domain Filtering

Because a well resolved tree is crucial for correct age assignment, we focus on eukaryotic genes. Contamination is a major problem in gene annotation (Kryukov & Imanishi 2016; Merchant et al. 2014; Salzberg 2017), so we filtered our dataset for likely contaminants, in addition to horizontally transferred genes. While horizontally transferred genes are not contaminants, their unusual form of inheritance means is likely to produce incorrect estimates of gene ages. The same applies to the unique evolutionary history of organelles.

Out of 17929 available pfams, we first excluded 4913 pfams that appeared in our eukaryotic dataset despite being annotated as occurring in Prokaryotes, but not Eukaryotes (as annotated in the tree file “pfamA_species_tree.txt”, accessed on May 17 2019 from https://pfam.xfam.org/).

Next we excluded pfams via keyword search. Of the 5076 pfams annotated as exclusive to Eukaryotes, 218 pfams that contained the terms “mitochondria” or “chloroplast” in the interproscan abstract were excluded, leaving 4858 pfams. Of these eukaryote-specific pfams, we further excluded 561 pfams for which there was no interpro abstract available, leaving 4297 pfams out of the 4858.

Of the 7773 pfams annotated as occurring both in Eukaryotes and in Bacteria, Archaea, or Viruses, we excluded 2064 contained the strings “viral”, “virus”, “bacter*”, “capsid”, “bacillus”, “Pilus”, “Pilin”, “mitochondria” or “chloroplast” unless they also contained one of the terms “eukary*”, “vertebrat”, “fung*”, “metazoa”, “plant”, “mammal”, “insect”, “yeast”, “human”, “all organisms”, “human”, “antibod”, “immune”, leaving 5709 pfams out of the 7773. Of these ancient pfams shared with prokaryotes, we further excluded 1112 pfams without an available interpro abstract, leaving 4597 pfams.

The 167 pfams that did not have species annotation information available in the pfam database (https://pfam.xfam.org/) were removed from the dataset, resulting in an overall total of 8894 pfams.

The phylogenetic distribution of some remaining pfams appeared indistinguishable from chance contamination. To formalize this criterion, we compared the number of losses inferred by Dollo parsimony to that occurring under the hypothesis that all hits are random contamination, performing this test for all pfams reported in less than half the total number of species. For each possible number of species containing the pfam, we simulated a distribution of the inferred number of losses when that number of species was chosen at random. We then used the mean and variance for the resulting distribution to calculate z-scores for the number of inferred losses from the actual data on each pfam, and rejected the pfam if z-scores > −2, a total of 688 pfams. This resulted in a final set of 8209 pfams.

We also remove all genes that contain a pfam excluded as described above.

### Age Assignment

#### Pfams

Domains were assigned a date of evolutionary origin using TimeTree. We assumed each pfam originated halfway between the node of the most recent common ancestor of all species in which the pfam originated, and the node prior to that. To add additional temporal resolution to the oldest domains, we identified any Pfams in our dataset likely to have been present in the first eukaryotic common ancestor, defined as those pfams present both in non-protist eukaryotes and in the protist group ‘Excavata’ (species included in our dataset are: *Naegleria gruberi, Giardia intestinalis, Giardia lamblia, Spironucleus salmonicida, Angomonas deanei, Bodo saltans, Leishmania braziliensis, Leishmania donovani, Leishmania infantum, Leishmania major, Leishmania mexicana, Leishmania panamensis, Leptomonas pyrrhocoris, Leptomonas seymouri, Perkinsela sp, Strigomonas culicis, Trypanosoma brucei, Trypanosoma cruzi, Trypanosoma rangeli, Trichomonas vaginalis*), which is considered a possible outgroup to the other eukaryotes, and with an estimated date of 2230 MYA (Hedges et al. 2001; Hedges et al. 2006; Parfrey et al. 2011).

Beyond TimeTree, pfams inferred to have been present in the last universal common ancestor (LUCA) (Weiss et al. 2016) were dated as 4090 MY old. For those pfams that are found in eukaryotes and bacteria, but not archaea, and also those pfams that are found in eukaryotes and archaea, but not bacteria, i.e. for those that emerged after LUCA but prior to the emergence of eukaryotes, we assigned them the age of 3145 MY, as the halfway point between the emergence of eukaryotes and LUCA. Dates of these ancient events are obviously imprecise.

#### Full Genes

We dated each gene according to the age of its oldest Pfam. Pfams represent highly conserved sequences, so it is unlikely that genes would be categorised as older based on non-Pfam sequences. Note that while different parts of a gene have different ages, here we classify a protein’s age based on the age of its oldest pfam.

#### Homology Groups

Homologous sequences share a common evolutionary origin, and therefore treating them as independent datapoints is a form of pseudoreplication (Wilson et al. 2017). To correct for this, an average taken across each Pfam was treated as a single datapoint in our domain analyses.

For genes, this was achieved by grouping according to their oldest Pfam. For genes with multiple Pfams that are all equally the oldest, a cluster analysis was performed. We started by considering the frequency of the concurrence of two Pfams, *A* and *B*, *P*(*AB*). If *P*(*AB*| *A or B*) ≥ 50% given all protein-coding genes in our dataset, then a link was established. Following all pairwise comparisons, Pfams were grouped together using single-link clustering, and each group was assigned a unique gene homology ID. An average across all genes sharing a gene homology ID was then treated as a single datapoint in our whole gene analyses.

The homology group files used in our analyses are available at doi:10.6084/m9.figshare.12037281.

### Metrics

#### Transmembrane Status

Genes and pfams were assigned as either transmembrane or non-transmembrane using the programme tmhmm (Krogh et al. 2001), which predicts the number and position of transmembrane helices within a protein sequence. A gene was designated as transmembrane if it contained over 18 amino acids that are predicted to be in a transmembrane helix (Krogh et al. 2001). A pfam was designated as transmembrane if it overlapped with a predicted transmembrane helix by a minimum of 50% of the pfam length, or if a transmembrane helix overlapped with a pfam by a minimum of 50% of the helix length. 50% was chosen as an arbitrary albeit reasonable cutoff for the designation of transmembrane status.

#### Amino Acid Composition

The fractional amino acid composition was found by counting the occurrences of each of the 20 amino acids in each protein or pfam and dividing by the length of the sequence. In order to obtain results with easily interpretable units, we show results for the untransformed proportions. Our qualitative results do not change if amino acid fractions are arcsine transformed prior to analysis (results not shown).

#### Disorder

Disorder predictions were made for each sequence using IUPred 2 (Dosztányi, Csizmok, et al. 2005; Mészáros et al. 2018), which assigns a score between 0 and 1 to each amino acid. Each protein’s ISD score was calculated by averaging over the values of all amino acids. To determine the disorder of each Pfam, we averaged over only the relevant subset. In order to obtain results with interpretable units, we show untransformed results, however, results are qualitatively similar if data is transformed (using a Box-Cox transform with the optimal value of lambda) prior to analysis. We also recalculated IUPred scores after first removing cysteine residues from protein sequences.

#### Clustering

To determine clustering scores, we followed Foy et al. (2019), and compared the variance of the hydrophobicity among blocks of *s*=6 consecutive amino acids to the mean hydrophobicity, to produce a normalized index of dispersion. If the length *L* of a protein was not a multiple of 6, we took the average of all possible *p* = (*L* modulo 6) frames, truncating the ends appropriately. The six most hydrophobic amino acids Leu, Ile, Val, Phe, Met and Trp were scored as +1, and the remaining standard amino acids to −1, as in Irback et al. (1996) and Foy et al. (2019). Non-standard amino acid residue abbreviations were also scored as +1 or −1, interpreted as follows: B as corresponding to D or N and hence −1; J as corresponding to either I or L and hence +1; Z as corresponding to either E or Q and hence −1; U as selenocysteine and O as pyrrolysine, both scored as −1.

For a sequence of length *N* = ⌊*L*/*s*⌋ in frame 1≤ *f* ≤ *p*, the scores for each transformed block *k* = 1,…,*N*/*s* of six amino acids were summed to σ_*k,f*_, with the full truncated sequence summing in total to

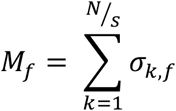

The normalized index of dispersion for a particular frame was then calculated as

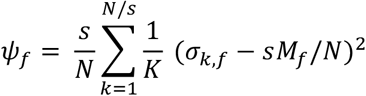

with the normalization factor

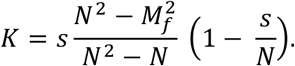

The total normalized index of dispersion was then calculated by taking the average over all possible frames

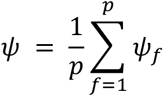

Clustering scores were again analyzed as raw scores for ease of interpretation of slope units, however, results are qualitatively similar if the data are transformed prior to analysis.

### Statistical Analyses

All statistical work in this research was performed using R. The function tidy from the broom package was used to convert model outputs to data frames for analysis and plotting, and to calculate 95% confidence intervals for regression slopes (Robinson 2014). We used the function lmer from the package lme4 (Bates et al. 2019) to perform linear mixed models and the function lm to perform basic linear models. The lm function was also used to perform weighted least squares regression on variances. Plots were generated in R, using packages ggplot2 and gridExtra. The parametric boxplots were drawn using the quantile function in base R.

### Data Availability

All scripts used in this work can be accessed at: https://github.com/MaselLab/ProteinEvolution. Our raw data, and homology files used in our analyses, are available at doi:10.6084/m9.figshare.12037281.

## Acknowledgements

This work was supported by the John Templeton Foundation (60814) and the National Institutes of Health (GM-104040).

## Supplementary Figures

**S1.**
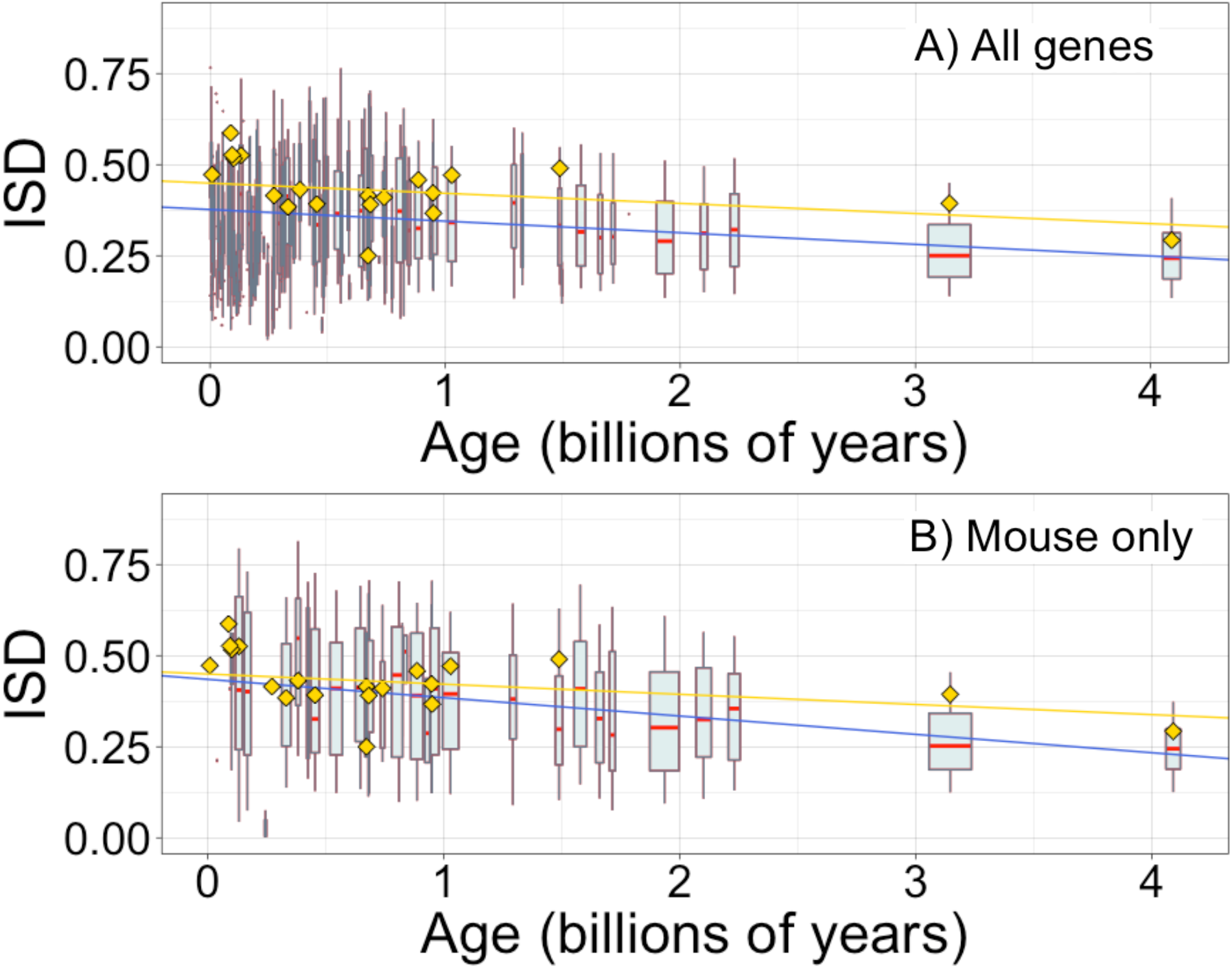
ISD depends on age for whole genes, as previously reported by Foy et al. (2019). Each data point consists of the average across all instances of a homologous gene family (see Methods) across all species (A) or just in mouse (B), dated according the oldest pfam the gene contains. Each age class is represented by a weighted box plot, where the width of the plot indicates the number of gene families in that age class. The median is shown in red, with the boxes representing upper and lower quartiles (the 75^th^ and 25^th^ percentile), and the whiskers indicating 9 and 91 quantiles. For age classes with only a single gene family, values are presented as small red dots. For clarity of presentation our plots do not show outliers. A) blue slope=−0.032, R^2^ = 0.057, p = 1×10^−229^. B) blue slope=−0.050, R^2^ = 0.091, p = 3×10^−85^. Slope represents the decrease in average IUPred2 score, i.e. the predicted propensity of the average amino acid to be disordered, per billion years. Yellow points show the average ISD scores and phylostratum assignments taken from Foy et al. (2019) and assigned dates using our scheme, with corresponding yellow linear regression with age (slope=−0.028, R^2^ = 0.042, p = 3×10^−136^). Our improved gene age assignments thus increased the strength of the relationship for mouse genes, where the relationship is stronger than for genes across all taxa.

**S2.**
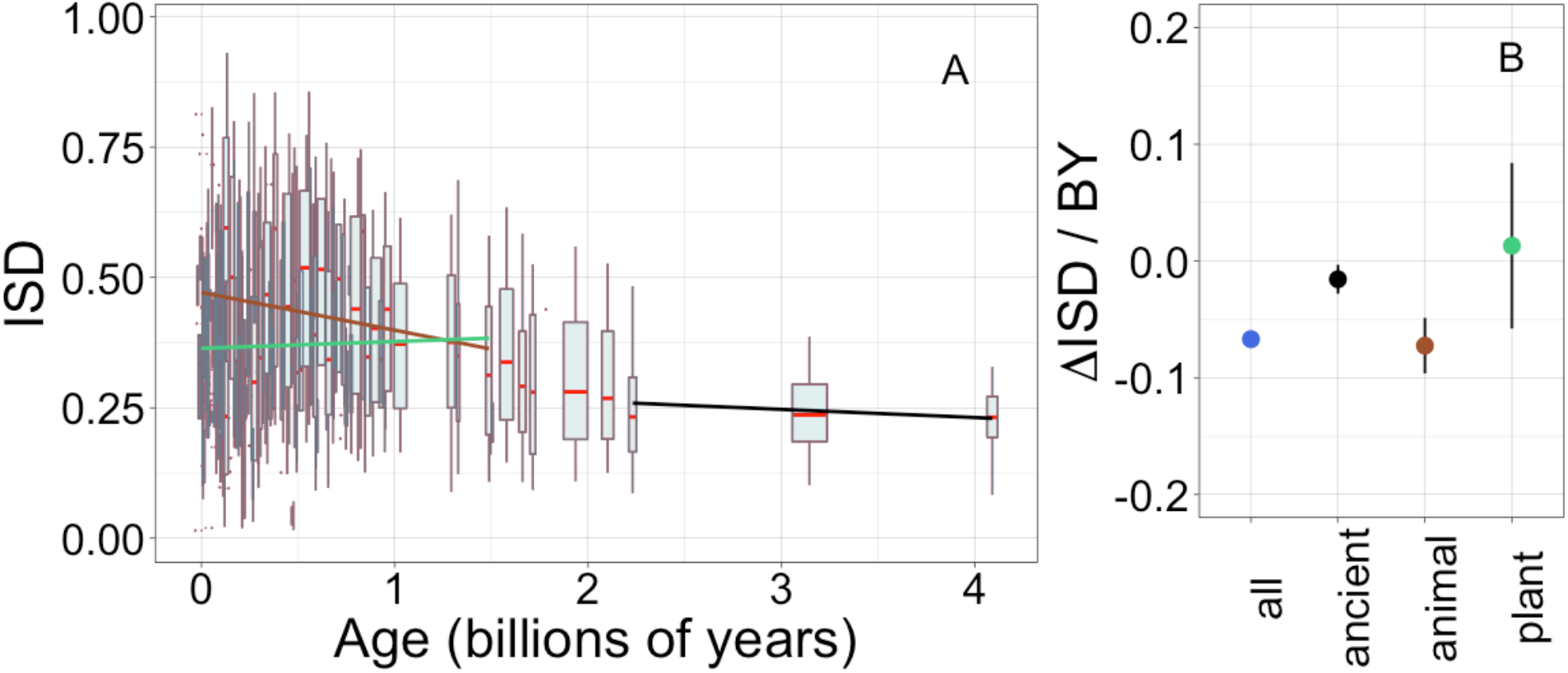
Recalculating ISD after excising cysteine residues has very little effect on our results. Details are as in figure 1. Regression calculated for recent animal pfams (brown slope=−0.073, R^2^ = 0.014, p = 3×10^−9^), recent plant pfams (green slope=0.013, p = 0.7), and ancient pfams in all lineages (black slope=−0.016, R^2^ = 0.0016, p = 0.014).

**S3.**
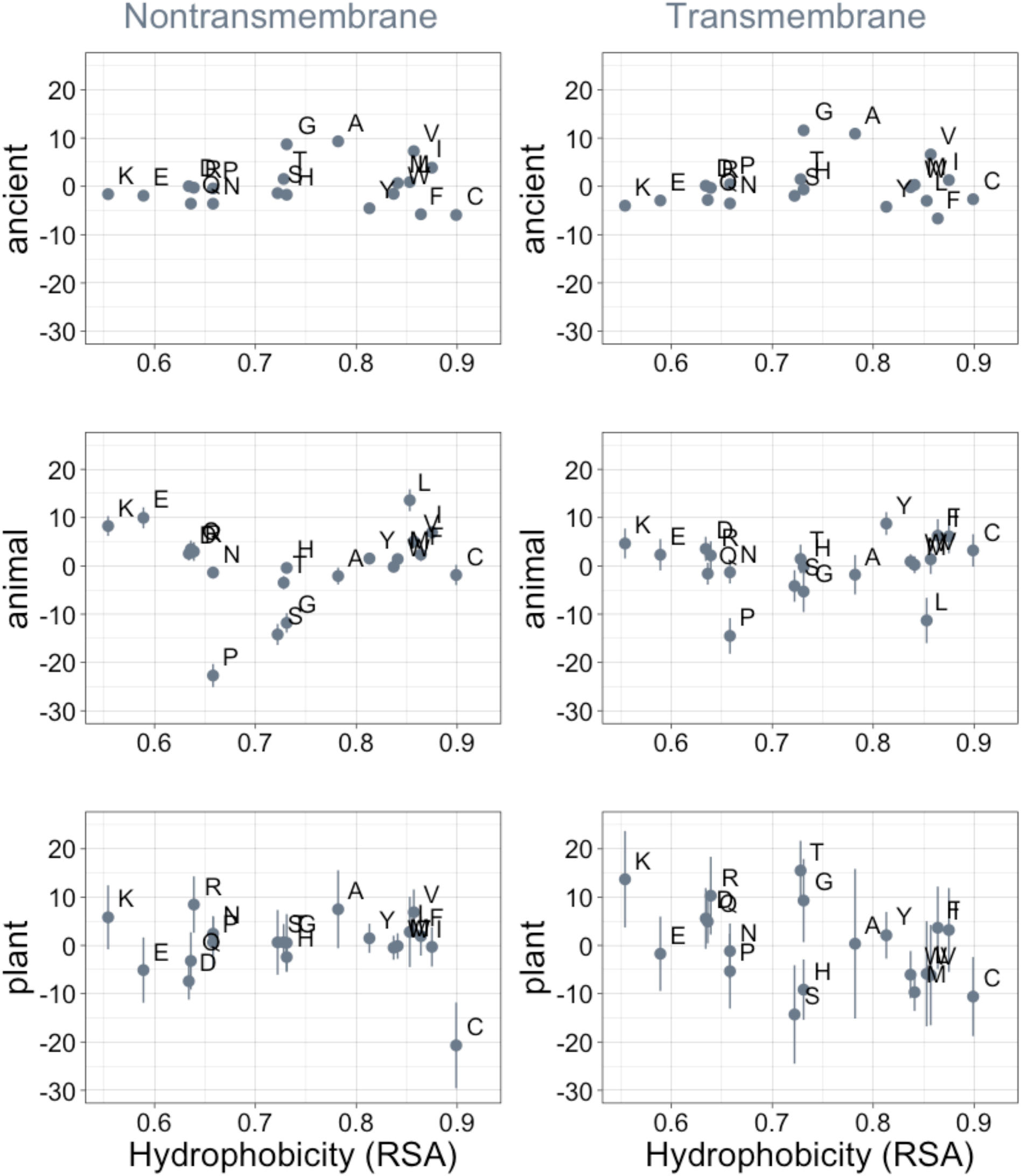
Phylostratigraphy slopes are not significantly correlated with hydrophobicity, as measured by 1-RSA (Tien et al. 2013). Phylostratigraphy slopes are in units of percentage points of composition per billion years. Lines indicate the standard errors on the slopes. ‘Ancient’ refers to pfams older than 2101 MY (nontransmembrane and transmembrane, Spearman’s p = 0.7 and 0.5), calculated over all lineages, whereas ‘animal’ (nontransmembrane and transmembrane, Spearman’s p = 0.8 and 0.6) and ‘plant’ (nontransmembrane and transmembrane, Spearman’s p = 0.9 and 0.05) slopes are calculated over pfams appearing after the divergence between the animal/fungi and plant lineages, 1496 MY, assessed in only animal or only plant instances.

**S4.**
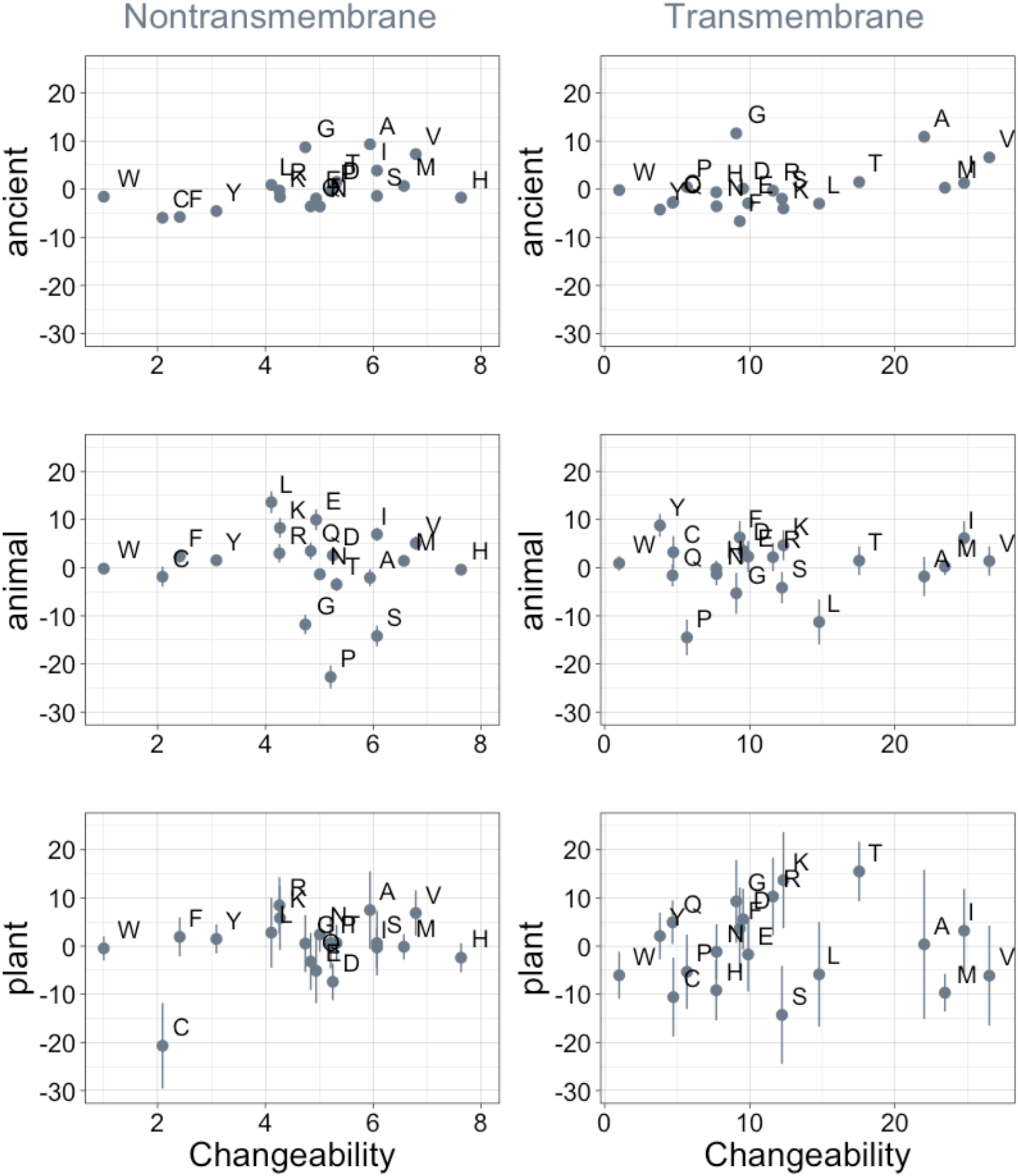
Phylostratigraphy slopes are not significantly correlated (after correction for multiple testing) with relative amino acid changeability. Changeability scores are relative to the least changeable amino acid, tryptophan (W), which is assigned a value of 1 in both the transmembrane and the non-transmembrane cases (Tourasse & Li 2000). Larger absolute changeability scores for transmembrane residues may therefore reflect low changeability of tryptophan, rather than higher changeability across the board. Phylostratigraphy slopes are in units of percentage points of composition per billion years. Insofar as changeability predicts phylostratigraphy slopes, especially for ancient trends, the relationship goes in the opposite direction to that predicted by homology detection bias. Lines indicate the standard errors on the slopes. ‘Ancient’ refers to pfams older than 2101 MY (nontransmembrane Spearman’s rho = 0.50, p = 0.02; transmembrane rho = 0.40, p = 0.08), calculated over all lineages, whereas ‘animal’ (nontransmembrane and transmembrane Spearman’s p = 0.53 and 0.9) and ‘plant’ (nontransmembrane and transmembrane Spearman’s p = 1.0 and 0.8) slopes are calculated over pfams appearing after the divergence between the animal/fungi and plant lineages, 1496 MY, assessed in only animal or only plant instances.

**S5.**
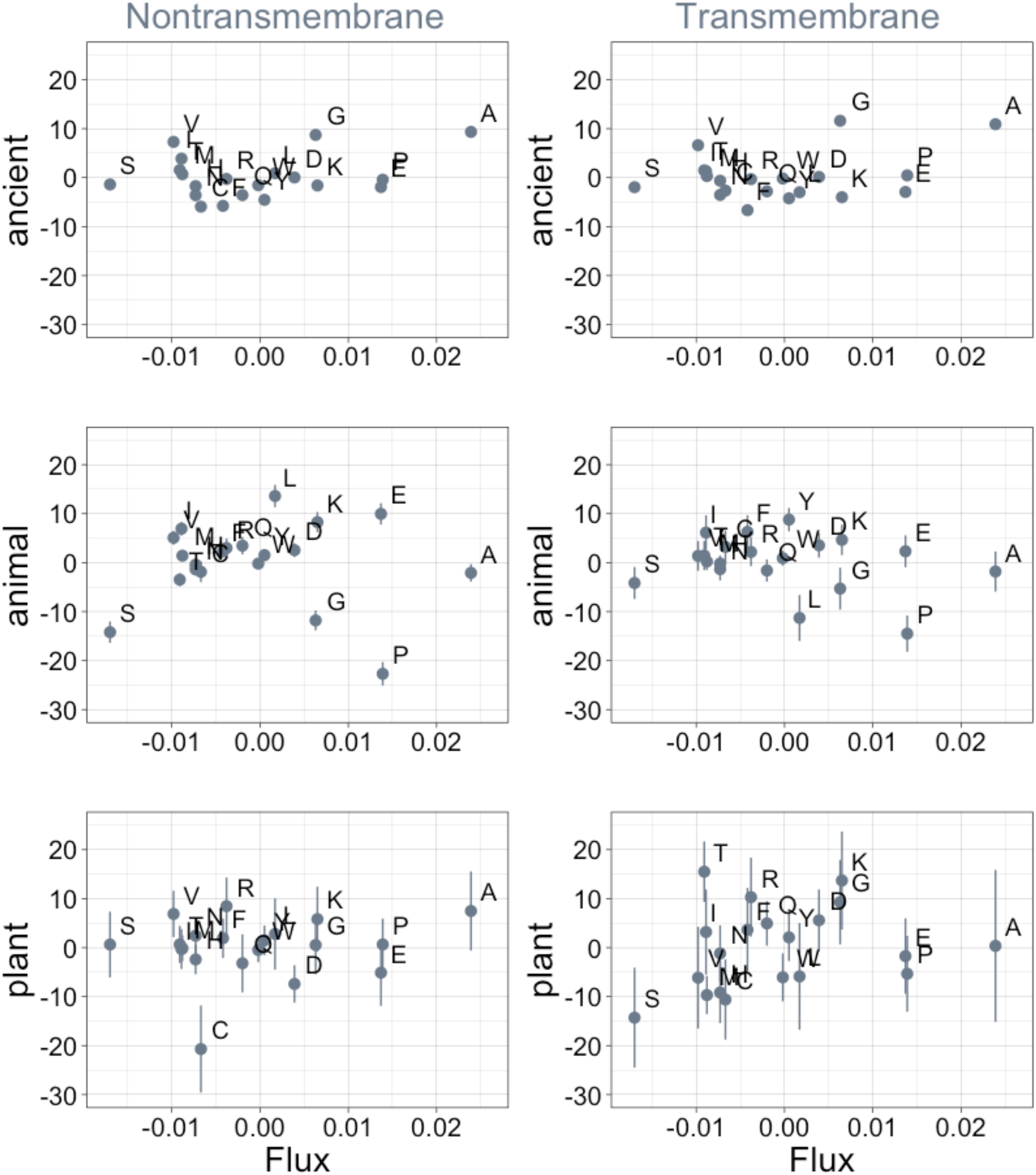
Phylostratigraphy slopes are not significantly correlated to the amino acid flux estimates of Jordan et al. (2005). Phylostratigraphy slopes are in units of percentage points of composition per billion years. Lines indicate the standard errors on the slopes. ‘Ancient’ refers to pfams older than 2101 MY (nontransmembrane and transmembrane, Spearman’s p = 1.0 and 0.7), calculated over all lineages, whereas ‘animal’ (nontransmembrane and transmembrane, Spearman’s p = 0.7 and 0.6) and ‘plant’ (nontransmembrane and transmembrane, Spearman’s p = 0.9 and 0.2) slopes are calculated over pfams appearing after the divergence between the animal/fungi and plant lineages, 1496 MY, assessed in only animal or only plant instances.

**S6.**
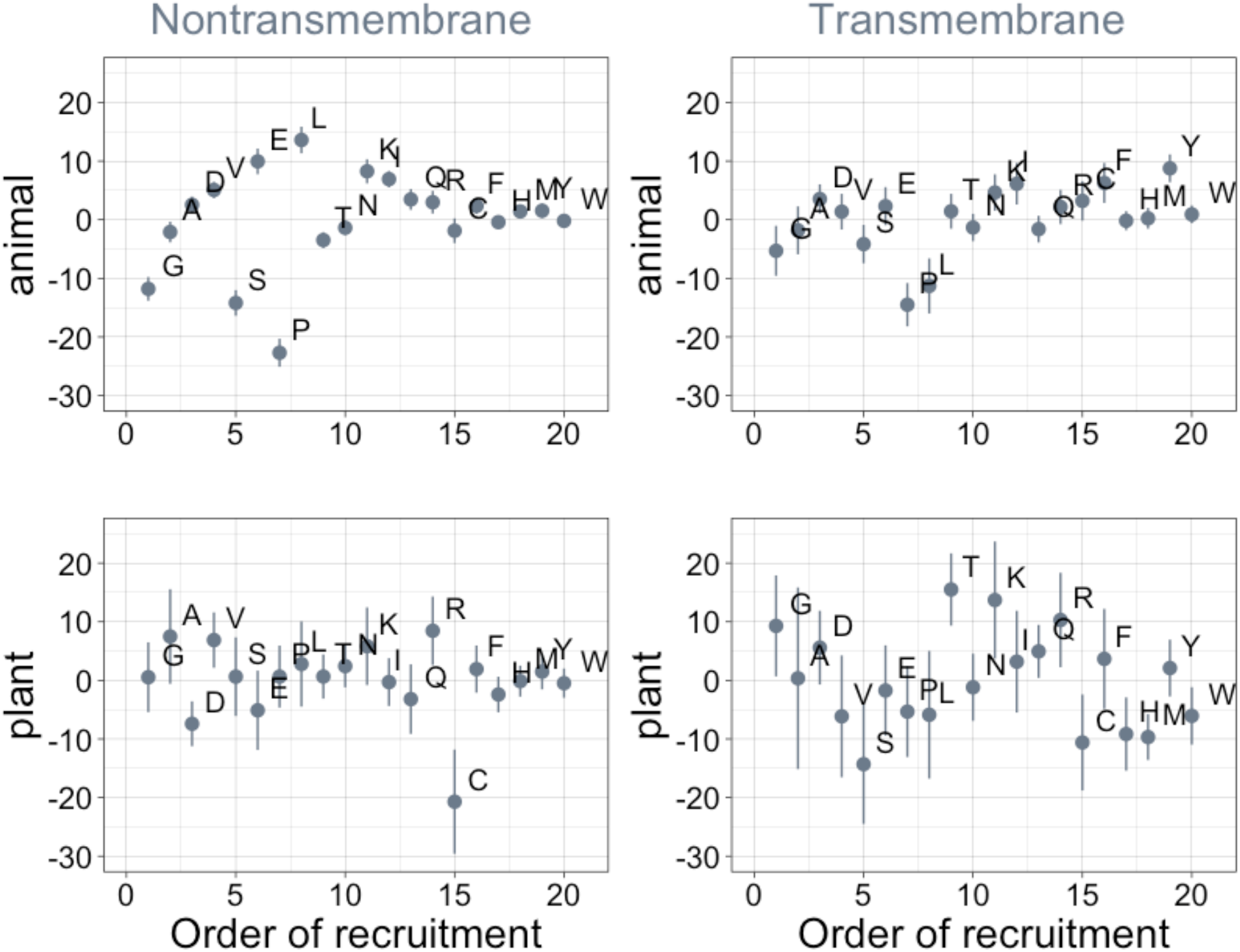
Order of amino acid recruitment does not affect domain composition in more recent lineages. Phylostratigraphy slopes are in units of percentage points of composition per billion years. Lines indicate the standard errors on the slopes. ‘Animal’ (nontransmembrane and transmembrane Spearman’s p = 0.6 and 0.07) and ‘plant’ (nontransmembrane and transmembrane Spearman’s p = 0.5 and 0.5) slopes are calculated over pfams appearing after the divergence between the animal/fungi and plant lineages, 1496 MY, assessed in only animal or only plant instances.

**S7.**
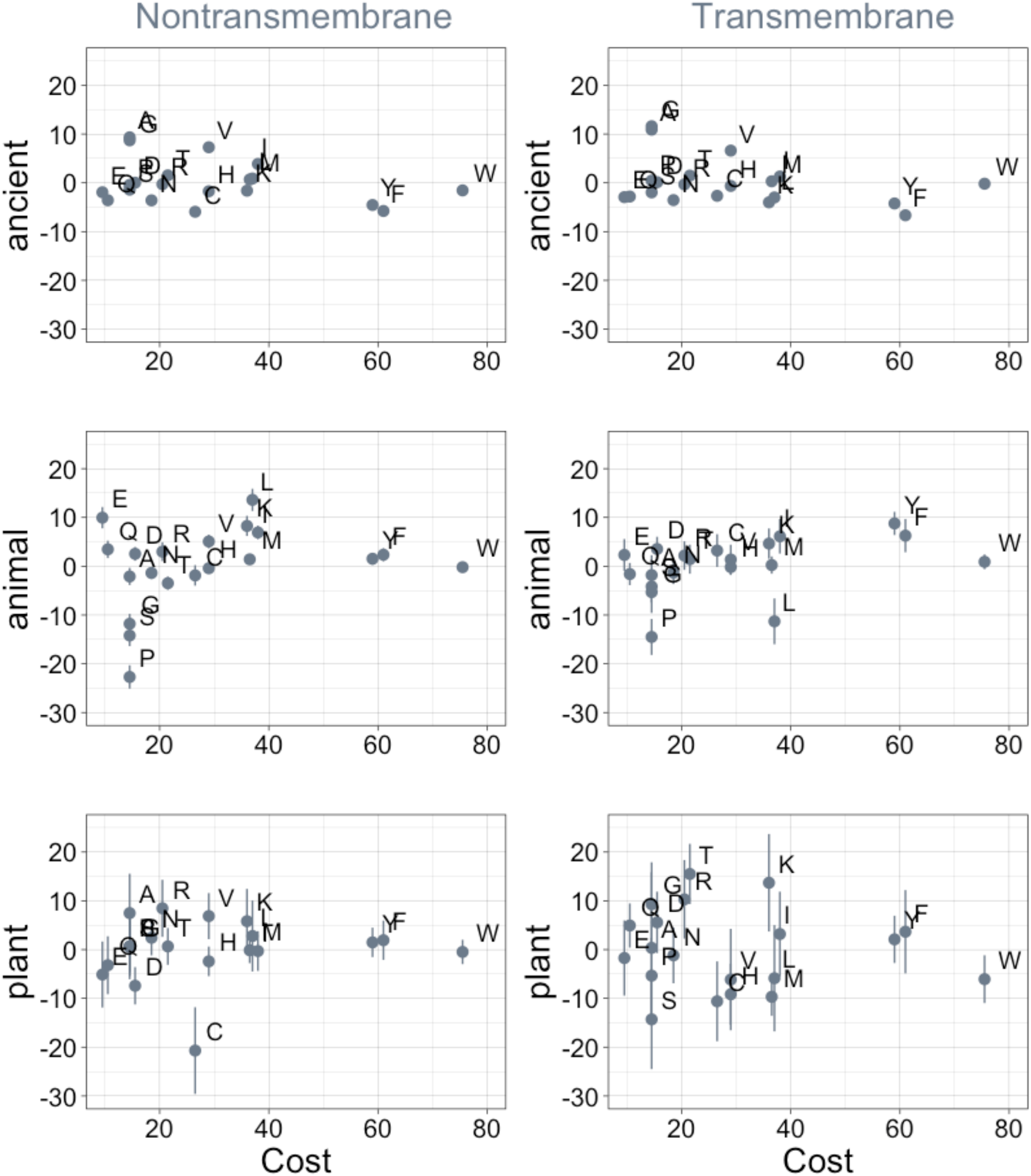
Phylostratigraphy slopes are not significantly correlated (after correction for multiple testing) to the cost of production of amino acids (aerobic metabolic cost, as estimated in yeast (Raiford et al. 2008)). Phylostratigraphy slopes are in units of percentage points of composition per billion years, with lines indicating the standard errors on the slopes. ‘Ancient’ refers to pfams older than 2101 MY (nontransmembrane and transmembrane, Spearman’s p = 0.6 and 0.3), calculated over all lineages, whereas ‘animal’ (nontransmembrane and transmembrane, Spearman’s p = 0.2 and 0.03) and ‘plant’ (nontransmembrane and transmembrane, Spearman’s p = 0.5 and 0.7) slopes are calculated over pfams appearing after the divergence between the animal/fungi and plant lineages, 1496 MY, assessed in only animal or only plant instances.

**S8.**
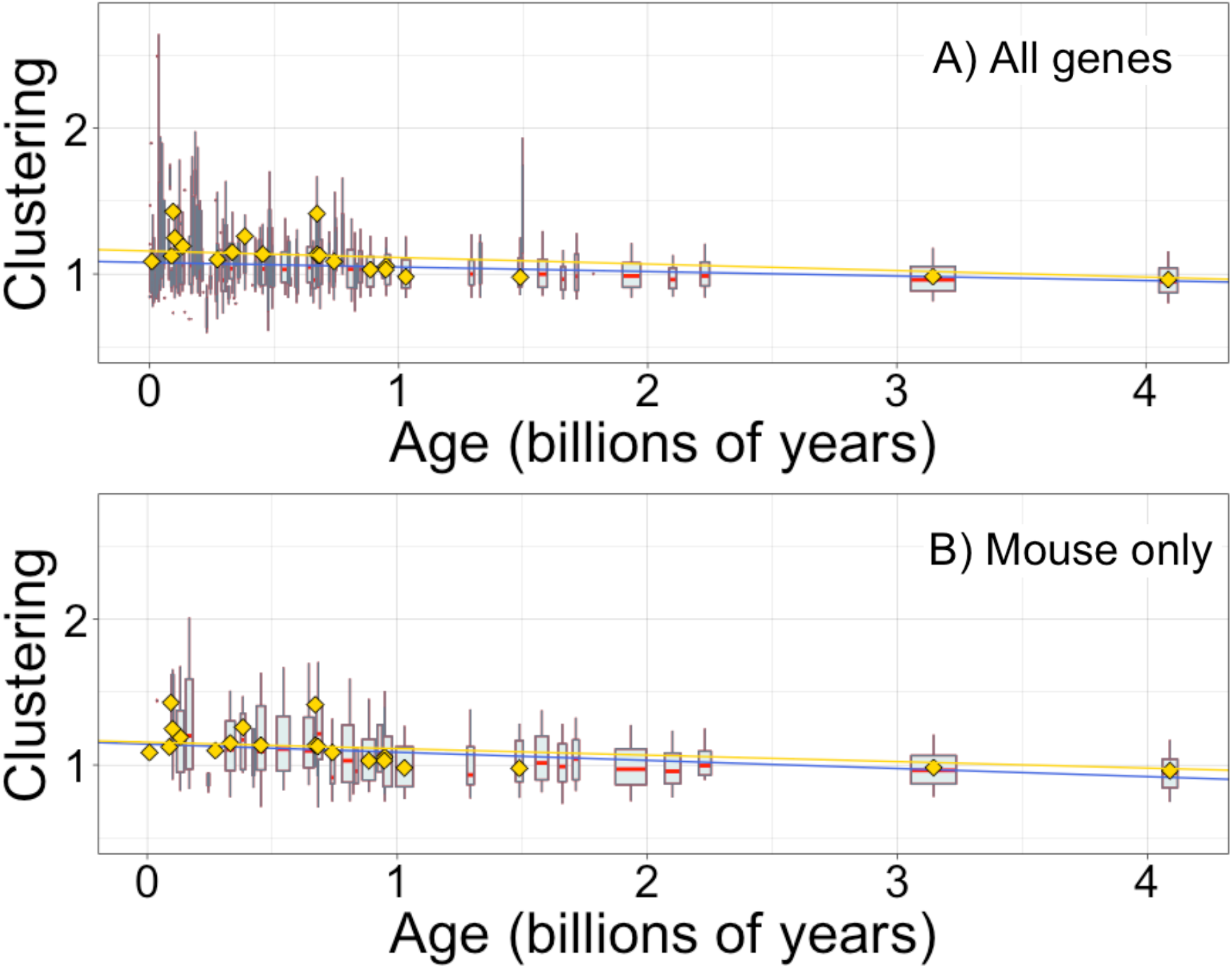
Hydrophobic clustering of complete genes depends on age, as previously reported by Foy et al. (2019). Clustering has an expected value of 1 for randomly distributed amino acids. Each data point consists of the average across all instances of homologous pfams, across all species in which it occurs. Phylostratigraphy assigns these to age classes, dated using timetree. Each age class is represented by a weighted box plot, where the width of the plot indicates the number of pfams in that age class. The median is shown in red, with the boxes representing upper and lower quartiles (the 75^th^ and 25^th^ percentile), and the whiskers indicating 9 and 91 quantiles. For age classes with only a single pfam, values are presented as small red dots. For clarity of presentation our plots do not show outliers. The relationship is a little weaker for all genes (A; blue linear regression slope=−0.031, R^2^ = 0.033, p = 5×10^−132^) than it is for genes found in mouse (B; blue linear regression slope=−0.056, R^2^ = 0.060, p = 4×10^−56^). In both plots, yellow points show the average clustering scores taken from Foy et al.’s (2019) mouse gene analysis, with corresponding yellow linear regression, slope= −0.045, R^2^ = 0.067, p = 8×10^−221^. Our improved gene age assignments thus increased the strength of the relationship for mouse genes, where the relationship is stronger than for genes across all taxa.

## Notes

https://github.com/MaselLab/ProteinEvolution

doi:10.6084/m9.figshare.12037281

